# Zoonotic viruses in three species of voles (*Microtus* spp.) from Poland

**DOI:** 10.1101/2020.08.22.259309

**Authors:** Maciej Grzybek, Katarzyna Tołkacz, Tarja Sironen, Sanna Mäki, Mohammed Alsarraf, Jolanta Behnke-Borowczyk, Beata Biernat, Joanna Nowicka, Antti Vaheri, Heikki Henttonen, Jerzy M. Behnke, Anna Bajer

**Affiliations:** Department of Tropical Parasitology, Medical University of Gdansk, Poland; Department of Parasitology, University of Warsaw, Warsaw, Poland; Department of Virology, University of Helsinki, Helsinki, Finland; Department of Forest Pathology, Poznan University of Life Sciences, Poznan, Poland; Natural Resources Institute Finland, Helsinki, Finland; School of Life Sciences, University of Nottingham, Nottingham, UK

**Author notes:** These authors contributed equally to this work (Maciej Grzybek, Katarzyna Tołkacz). These authors contributed equally to this work (Jerzy M. Behnke, Anna Bajer). **ADDRESS FOR CORRESPONDENCE:** Maciej Grzybek, PhD; Department of Tropical Parasitology, Medical University of Gdansk, Powstania Styczniowego 9B, 81-519 Gdynia, Poland.

**Keywords:** Arenavirus, hantavirus, LCMV, CPXV, PUUV, TULV, *Microtus*, vole, seroprevalence, Poland

## Abstract

Rodents are known to be reservoir hosts for a plethora of zoonotic viruses and therefore play a significant role in the dissemination of these pathogens. We trapped three vole species (*Microtus arvalis, M. oeconomus* and *M. agrestis*) in N.E. Poland, all of which are widely distributed species in Europe, and, using immunofluorescence assays, we assessed serum samples for the presence of antibodies to hantaviruses, arenaviruses and cowpox viruses (CPXV). We detected antibodies against CPXV and Puumala virus (PUUV), overall seroprevalence of combined viral infections of 18.2% [10.5-29.3] and mostly attributed to CPXV. Antibodies to Tula hantavirus, typical for *Microtus* voles, are known to cross-react strongly with the PUUV antigen used in PUUV screening, but we detected only one PUUV/TULV cross-reaction in *Microtus arvalis* (1.3% [0.1-7.9]). We found antibodies against CPXV in all three *Microtus* spp. Seroprevalence was similar in all vole species. There were no significant differences in seroprevalence between the sexes and among host age categories. Our results contribute to the increasing understanding of CPXV abundance in voles in Europe, and confirm that CPXV circulates also in *Microtus* spp. voles in NE Poland.

## 1. INTRODUCTION

The identification of possible hosts, and the study of transmission dynamics in their populations, are both crucial steps in controlling zoonotic diseases (Luis et al., 2013; Mills, 1998). Rodents, as the most widespread and abundant mammals are considered to be among the most significant source of zoonotic pathogens (Bordes et al., 2015; Luis et al., 2013). Studies of population dynamics have demonstrated that European rodent populations experience multiannual and cyclic density fluctuations (Cornulier et al., 2013; Hansson and Henttonen, 1985; Sokolik et al., 2015). This has been linked with variation in the incidence of zoonoses spread by rodents (Kallio et al., 2009; Olsson et al., 2009).

Rodent-borne hanta-, arena- and orthopox viral pathogens are maintained in nature by direct intraspecies or, or as probably is the case with cowpox virus (CPXV), interspecies transmission from rodent to rodent without the participation of arthropod vectors. Viral infection is usually persistent in hantaviruses, possibly also in arenaviruses, but in CPXV viremia lasts for only 2-3 weeks. Transmission in rodents occurs by contact with body fluids or excretions (Carroll et al., 2015; Charrel and de Lamballerie, 2010; Kallio et al. 2007, Vapalahti et al., 2003). The most prevalent rodent-borne viruses carried by European rodents include hantaviruses, lymphocytic choriomeningitis virus (LCMV), CPXV (Goeijenbier et al., 2013; Grzybek et al., 2019; Jääskeläinen et al., 2013; Kallio-Kokko et al., 2005; Kinnunen et al., 2011; Tonteri et al., 2013). Puumala virus (PUUV) is widely prevalent in bank vole (*Myodes glareolus*) populations (Brummer-Korvenkontio et al., 1982; Vapalahti et al., 2003). Hantavirus infections in bank voles are chronic and therefore viral replication and shedding is persistent (Kallio et al., 2007; Voutilainen et al., 2015). As a consequence, the rodent host may be infectious for the whole of its lifespan (Meyer and Schmaljohn, 2000).

CPXV is the only known wildlife-borne Orthopox virus (OPV) in Europe (Kinnunen et al., 2011). Field voles (*Microtus agrestis*), among other rodent species (i.e. *Apodemus* sylvaticus and *Myodes glareolus*), are known to act as reservoir hosts for CPXV (Chantrey et al., 1999; Crouch et al., 1995; Kinnunen et al., 2011). LCMV, the only arenavirus in Europe, was thought to be prevalent in the house mouse (*Mus musculus*) (Blasdell et al., 2008). However, further studies revealed that the virus is present in other murine and vole species (Blasdell et al., 2008; Kallio et al., 2006; Laakkonen et al., 2006; Tagliapietra et al., 2009). Ledesma et al. (2009) identified an independent genetic lineage of LCMV in wood mice and this led to the suggestion of spillover and the circulation of multiple related and cross-reactive Mammarena viruses.

Voles (*Microtus* spp. and *Myodes* spp.) are the most abundant rodents in European grasslands and forests (Hanski and Henttonen, 1996). The occurrence of rodent-borne viruses is not well recognized, nor indeed documented in Poland. Therefore, this study aimed to evaluate zoonotic and potentially zoonotic viruses in a population of *Microtus* spp. in NE Poland. Our results make a significant contribution to the understanding of the role of vole populations in the maintenance and dissemination of these viral pathogens in our geographical region.

## 2. MATERIAL AND METHODS

### 2.1. Ethical approval

This study was carried out with due regard for the principles required by the European Union and the Polish Law on Animal Protection. Formal permits were obtained, allowing trapping in the field and subsequent laboratory analysis of sampled materials. Our project was approved by the First Warsaw Local Ethics Committee for Animal Experimentation (ethical license numbers: 148/2011 and 406/2013).

### 2.2. Collection of voles

The study site was located in the Mazury Lake District region in the north-eastern corner of Poland (Urwitałt, near Mikołajki; 53°48’50.25”N, 21°39’7.17”E) and previously described (Tołkacz et al., 2018, 2017). Voles were collected in August 2013 during the late summer season, when rodent population density is at its highest in the annual cycle. Voles were live-trapped using mixed bait comprising fruit (apple), vegetables (carrot and cucumber) and grain. Two traps were set every 10 m along the trap lines at dusk. The following morning traps were checked and closed to prevent animals from entering during daytime and to avoid losses from excessive heat from exposure of traps to direct sunlight. Traps were then re-baited and reset on the following afternoon. All traps were also closed during periods of intensive rainfall. All captured voles were transported in their traps to the laboratory for inspection.

The autopsies were carried out under terminal isoflurane anaesthesia. Animals were weighed to the nearest gram, total body length and tail length were measured in millimetres. Animals were allocated to three age classes (juveniles, subadult and adults), based on body weight and nose-to-anus length together with reproductive condition (scrotal, semi-scrotal or non-scrotal for males; lactating, pregnant or receptive for females) (Haukisalmi et al., 1988; Pawelczyk et al., 2004; Tołkacz et al., 2018, 2017). We confirmed the species identity by examination of the lower molars M_1_ and M_2_ and the second upper molar M^2^, especially to distinguish between juvenile individuals of *M. oeconomus* and *M. agrestis* (Pucek, 1981).

Blood samples were collected directly from the heart using a sterile 1.5 mL syringe immediately after death from over-exposure to Isoflurane (Baxter, USA) anaesthetic. Samples were centrifuged at 5,000 rpm for 10 min using an MPW High-Speed Brushless Centrifuge. Serum was collected and stored at −80°C until the samples could be analysed on completion of the fieldwork.

### 2.3. Serological screening of virus antibodies

*Microtus* spp. serum samples were analysed using an immunofluorescence assay (IFA). The serum samples were diluted 1:10 in PBS and the reactivity of the samples to hantaviruses was tested with PUUV-(Puumala virus)-IFA, to cowpox viruses with CPXV (Cowpox virus)-IFA and arenaviruses with LCMV (Lymphocytic choriomeningitis virus)-IFA. PUUV (Sotkamo strain), CPXV, and LCMV (Armstrong strain) -infected Vero E6 cells were detached with trypsin, mixed with uninfected Vero E6 cells (in a ratio of 1 : 3), washed with PBS, spotted on IFA slides, air-dried, and fixed with acetone as described earlier (Hedman et al., 1991). The slides were stored at −70°C until use. TULA orthohantavirus (TULV), specific for *Microtus* voles, is known to cross react strongly with PUUV antibodies (and *vice versa*). Thus, we report it as PUUV/TULV seroprevalent.

IFAs were carried out as previously described (Kallio-Kokko et al., 2006) with seropositive human serum as a positive control for the PUUV- and CPXV-IFA; and LCMV mouse monoclonal antibody (Progen, Heidelberg) for the LCMV-IFA. The slides were read under a fluorescence microscope and photographs were taken with a ZOE™ fluorescent cell imager (BioRad).

### 2.4. Statistical analysis

Prevalence values (percentage of animals infected) are given with 95% confidence limits in parenthesis (CL_95_) or error bars on figs, calculated by bespoke software based on the tables of Rohlf and Sokal (Sokal and Rohlf, 1994).

The statistical approach has been documented comprehensively in our earlier publications (Bajer et al., 2005; Behnke et al., 2008, 2001; Grzybek et al., 2018, 2015). For analysis of prevalence, we used maximum likelihood techniques based on log-linear analysis of contingency tables in the software package IBM SPSS Statistics Version 21 (IBM Corporation).

## 3. RESULTS

In total, we screened 77 *Microtus* spp. serum samples for the presence of arenavirus and hantavirus antibodies. We confirmed the presence of antibodies against CPXV and PUUV/TULV. No individuals were seropositive to LCMV.

The overall seroprevalence of zoonotic viruses was 18.2% (10.5-29.3). However, most of the inspected individuals were seropositive for CPXV (16.9% [9.4-27.9]) with only one individual showing evidence for presence of hantaviral antigens (1.3% [0.1-7.9]). Further analysis is therefore confined to seroprevalence of CPXV.

CPXV antibodies were present in all three *Microtus* spp. However, there were no significant differences between vole species (Figure 1A). Female voles had a marginally higher value for CPXV seroprevalence than males (18.2%[8.4-34.8] and 15.2%[7.0-28.7], respectively) (Figure 1B), but the difference between the sexes was not significant. There was no significant effect of host age on CPXV seroprevalence (Figure 1C).

**Figure 1.**
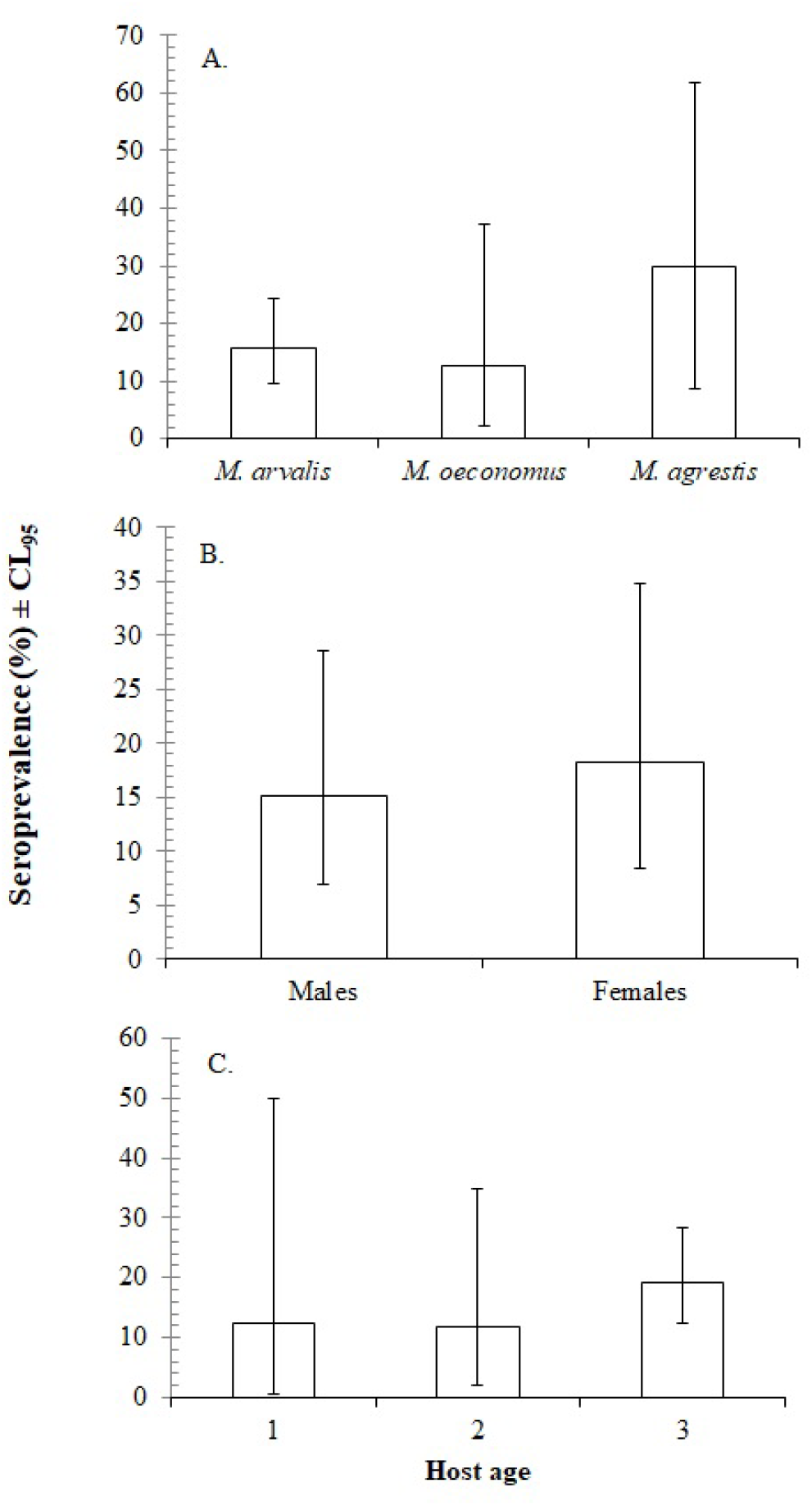
Seroprevalence of CPXV within: (A) three *Microtus* spp; (B) host sex; (C) host age.

## 4. DISCUSSION

Our results confirm that voles do host viral pathogens and hence are likely to play a role as reservoirs of viral infections with zoonotic potential, i.e. that constitute a significant threat to human and livestock health. We report a high overall seroprevalence of viral infections (18.2%). These findings are not only of considerable relevance to public health in the region but by inference are likely to have relevance also for other European regions populated by *Microtus* spp.

In our data the overall seroprevalence of viral infections is mostly accounted for by CPXV. This is consistent with our report on seroprevalence of zoonotic viruses in bank voles (*M. glareolus*) from the same geographical region (Grzybek et al., 2019). It is also in agreement with those obtained in different parts of Europe (e.g. Finland, England and Turkey) where CPXV, PUUV and TULV virus species have been detected in voles (Forbes et al., 2014; Laakkonen et al., 2006; Razzauti et al., 2009; Schmidt-Chanasit et al., 2010; Thomason et al., 2017). We did not detect any seropositivity to LCMV in the *Microtus* spp. individuals that we sampled (but see Forbes et al., 2014).

The current work was based on the presence/absence of specific antibody against viruses, and hence positivity in our assay reflected the history of previous infections and not necessarily current infection (Grzybek et al., 2018). Intrinsic factors such as host age, maturity and host sex may all influence the host’s exposure to viral infections and hence seroprevalence of antibodies to the specific pathogens that may have been experienced by animals (Grzybek et al., 2019, 2015). On this basis we had expected to find a higher seroprevalence among the older animals which would have had more opportunity for exposure to, and hence experience of infection than juveniles, but the difference between the age classes in the current study was not significant..

In summary, using cross-reacting PUUV antigen, we found serological evidence for the presence of CPXV and TULV in *Microtus* vole populations in NE Poland. We believe that identifying rodent species that can serve as reservoirs of zoonotic diseases and predicting regions where new outbreaks are most likely to happen, are crucial steps in preventing and minimising the extent of zoonotic diseases in humans (Han et al., 2015). Our results help to consolidate the gap in knowledge about the role of Polish voles as possible candidates for reservoirs of viral infections.

## ACKNOWLEDGEMENTS

We thank the University of Nottingham, Warsaw University and the Medical University of Gdansk for financial support. We appreciate the support from Sigrid Jusélius Foundation, Helsinki and Natural Resources Institute Finland. MG was supported by the National Science Centre, Poland under BiodivERsA3 programme (2019/31/Z/NZ8/04028) and by the Polish National Agency for Academic Exchange under the Bekker programme (PPN/BEK/2019/1/00337). MG thanks Alicja Rost and Ewa Zieliniewicz for their assistance in laboratory.

## AUTHORSHIP

The study was conceived and designed by MG, TS, KT and BB. Samples were collected in the field by KT, MA, JBB, JMB, AB, MG. The immunological analysis and laboratory work was conducted by MG, SM, JN, AV and TS. Data analysis was carried by MG & JMB. The ms was written by MG & KT, in consultation with all co-authors. MG, JMB, AB, HH revised the manuscript. All authors accepted the final manuscript version.

